# Optimised proteomic analysis of insulin granules from MIN6 β-cells identifies Scamp3, a novel regulator of insulin secretion and content

**DOI:** 10.1101/2024.04.23.590838

**Authors:** Nicholas Norris, Belinda Yau, Carlo Famularo, Helen E. Thomas, Mark Larance, Alistair M. Senior, Melkam A. Kebede

## Abstract

Pancreatic β-cells in the islets of Langerhans are key to maintaining glucose homeostasis, by secreting the peptide hormone insulin. Insulin is packaged within vesicles named insulin secretory granules (ISGs), that have recently been considered to have intrinsic structures and proteins that regulate insulin granule maturation, trafficking, and secretion. Previously, studies have identified a handful of novel ISG-associated proteins using different separation techniques. Here, this study combines an optimized ISG isolation technique and mass spectrometry-based proteomics, with an unbiased protein correlation profiling and targeted machine learning approach to uncover 211 ISG-associated proteins. Five of these proteins: Syntaxin-7, Synaptophysin, Synaptotagmin-13, Zinc transporter ZIP8 and SCAMP3 have not been previously ISG-associated. Through colocalization analysis of confocal imaging we validate the association of these proteins to the ISG in MIN6 and human β-cells. We further validate the role for one (SCAMP3) in regulating insulin storage and secretion from β-cells for the first time. SCAMP3 knock-down INS-1 cells show a reduction in insulin content and dysfunctional insulin secretion. These data provide the basis for future investigation into β-cell biology and the regulation of insulin secretion.

**Article Highlights:**

a. Why did we undertake this study? We undertook this study to optimize insulin granule isolation techniques alongside enhanced proteomics analyses to establish the first published murine insulin granule proteome.
b. What is the specific question(s) we wanted to answer? We aimed to specifically answer and investigate what proteins are present on insulin granules from MIN6 cells to further our understanding of insulin granule biogenesis, trafficking, and secretion.
c. What did we find? We find and validate the presence of 5 novel insulin granule-associated proteins.
d. What are the implications of our findings? An extensive proteomics analysis of MIN6 insulin granules and implicate Scamp3 as a novel protein that regulates insulin content and secretion in beta-cells.

## INTRODUCTION

Pancreatic β-cells play a crucial role in maintaining blood glucose levels and their dysfunction is a hallmark of type 2 diabetes. Located within the I*slets of Langerhans,* these cells release insulin, which stimulates glucose uptake by adipose tissue, liver, and muscle^1^. In type 2 diabetes, β-cells exhibit a reduction in glucose-stimulated insulin secretion (GSIS), and dysregulated insulin biogenesis and trafficking^2,3^.

Within the pancreatic β-cell, insulin is stored within the insulin secretory granules (ISGs), which are ∼300 nm in diameter and contain a dense core^4^. In response to nutrient stimuli ISGs are triggered for exocytosis, transported to the plasma membrane where they fuse and release insulin.

ISG biogenesis begins at the trans-Golgi network (TGN), where proinsulin is sorted alongside prohormone convertases, other secreted hormones and carrier proteins^4^ into immature ISGs. Within these immature ISGs, prohormones are cleaved into mature forms^5^, granule lumen become acidified and undesired proteins are removed^6^. Mature insulin then crystallizes with zinc cations to form insulin hexamers, constituting the dense core of the ISGs^7^.

ISGs are dynamic structures, and numerous proteins residing within them play crucial roles in insulin biogenesis and secretion, with implications for type 2 diabetes. For example, deficiencies in Chromogranin-B (CHGB) leads to impaired proinsulin processing, reduction in GSIS *in vivo*^8^ and *in vitro*^9^. Knock-out models of protein interacting with C Kinase-1 (PICK1) and islet cell autoantigen of 69 kDa (ICA1) also cause impaired insulin secretion and defective glucose metabolism^10^. Dysfunction of these ISG resident proteins contribute to metabolic disease and type 2 diabetes. Thus, interrogating the ISG and discovering novel ISG-resident proteins will enhance our understanding of insulin granule biogenesis, trafficking, and glucose-mediated secretion of ISGs.

Four previous studies have attempted to capture snapshot proteomes of ISG within β-cells^11–14^. Typically, distinct organelle proteomes are obtained through subcellular fractionation methods and fluorescent-assisted organelle sorting (FAOS), using tagged fluorescent organelle markers. However, achieving an accurate ISG proteome is challenging due to their dynamic nature and association with many other organelles within the β-cell^14^. Notably, the above-mentioned studies only assemble 5 proteins in consensus: Insulin-1 (Ins1), Insulin-2 (Ins2), Chromogranin-A (CgA), Prohormone convertase 2 (PC2) and Carboxypeptidase E (CPE). Given this, it is difficult to interpret the collective proteomic data due to the co-enrichment of contaminating proteins or fragments of subcellular organelles such as the Golgi network, lysosomes and most notably and commonly, mitochondria^11–14^. Our study addresses this caveat by employing an optimized protocol to enrich for ISGs and a dual unbiased and targeted proteomics approach to minimize contamination from other subcellular compartments.

Previous studied on ISG isolation methods and proteomics analyses have focused solely on rat insulinoma INS-1 or INS1-E cell lines^11–14^. In this study, we modify an ISG enrichment method described by Chen et al. 2015 ^15^, to enrich ISGs from the mouse insulinoma β-cell line, MIN6. MIN6 cells are arguably a better model for human metabolism because they possess a larger pool of insulin granules, respond similarly to human islets in response to lipotoxicity^16^ and can form pseudo-islets in culture^17^. The modified three-step purification protocol involves sequential Optiprep and Percoll gradient enrichment followed by sucrose purification of mature ISGs. LC-MS/MS analysis of fractionation samples identified 8021 unique peptides. Using protein correlation profiling (PCP) analysis, we identify 432 potential ISG proteins. Parallel analysis using proteomic-specific software allowed us to categorize proteins into defined subcellular organelles in MIN6 cells, assigning 211 proteins to the ISG with high confidence. To validate ISG targets, we conducted insulin colocalization analysis in immuno-stained MIN6 cells and human islets, confirming the ISG localization of SCAMP3, Synaptophysin, Synaptotagmin-13, Syntaxin-7 and ZIP8. Finally, siRNA-mediated SCAMP3 knockdown in INS-1 cells elicited a defect in insulin storage, and GSIS, implicating the novel ISG protein in healthy β-cell insulin regulation for the first time.

## Material and Methods

### Human pancreata and ethics statement

Human pancreata tissue were obtained from deceased organ donors as a part of the St Vincent’s Institute islet isolation program, through Donatelife to obtain research consent and adhere to strict ethical guidelines developed by the National Health and Medical Research Council (NHMRC), Australia (Reference Number: E76, ISBN: 9781925129595). Donor information including HbA1c, diabetes status, age, BMI, and sex can be found in Supplementary Table 5.

### Cell Culture

The MIN6 mouse insulinoma β-cells were purchased from AddexBio^TM^ and cultured in standard culture media as stated previously^18^. INS-1 cells were cultured in standard cultured media as previously described^18^. Culture media was changed every 2 – 3 days and cells were passaged every 4 – 5 days. MIN6 cell passages used were between 15 and 26. INS-1 cell passages used were between 19-33.

### Optimized three-step purification of mature ISGs from MIN6 cells

In this study, a two-step subcellular fractionation protocol described in a previous study Chen et al. 2015 was optimized. MIN6 cells were grown on 15 cm diameter petri dishes to 80 – 90% confluence. Plates were PBS washed, then cells scraped into 2 mL of ice-cold PBS and centrifuged at 300 x g for 3 min. The pellet was resuspended in 1 mL of Buffer A (0.3M sucrose, 1 mM EDTA, 1 mM MgSO_4_, 10 mM MES-KOH, pH 6.5 containing EDTA-free protease inhibitors (Roche), then passed through a 21 G and 25 G gauge needle (10 times each). Homogenates were centrifuged (1000 x g, 5 min) and the supernatant was loaded atop a discontinuous Optiprep gradient of five 2mL layers: (from top): 8.8, 13.2, 17.6, 23.4 and 30 %, diluted in Buffer B (2 mM EGTA, 20 mM MES-KOH, pH 6.5). Samples were centrifuged (100,000 x g, 80 mins) in a P50AT2 fixed angle rotor in a Hitachi 100NX ultracentrifuge set at half acceleration to ensure Optiprep layers were undisturbed, and half deceleration. 1 mL fractions were removed from the top to bottom of the tube to obtain a total of 16 fractions. Percoll was used in the second step to further enrich and purify ISGs. Mature ISG-enriched Fractions 11 and 12 were loaded on 10 mL of 27% Percoll solution, diluted in Buffer A in a 16 mL polypropylene tube, then centrifuged (35,000 x g, 60 min) in a A22-24×16 (Thermo Fisher) fixed-angle rotor in a Sorvall Lynx 4000 Centrifuge (Thermo Fisher). 1 mL fractions were removed from the top to bottom to obtain 12 fractions. In the third step, twelve 1 mL fractions following Percoll enrichment were suspended in 9 mL each of sucrose wash buffer (0.3 M sucrose, 5 mM HEPES, 1 mM EGTA, pH 7) and centrifuged (23,700 × g, 15 min) to pellet ISGs and remove Percoll from solution. Supernatants were discarded and the pellet was resuspended in 4% sodium deoxycholate (SDC) in water and boiled (100 °C, 10 min) for MS analysis.

### LC-MS/MS

LC-MS/MS samples were prepared as previously described^19^. Peptides were reconstituted with 5% formic acid in MS-grade H2O, sealed and stored at 4°C until LC-MS/MS acquisition. Using a ThermoFisher RSLCnano ultrahigh performance liquid chromatograph, peptides in 5% (v/v) formic acid (3 µL injection volume) were injected onto a 50 cm x 75 µm C18Aq (Dr Maisch; 1.9 µm) fused analytical column with a ∼10 µm pulled tip, coupled online to a nanospray electrospray ionization source. Peptides were resolved over gradient from 5 – 40 % acetonitrile for 70 min, with a flow rate of 300 nL min^-1^ (Capillary flow). Electrospray ionization was done at 2.3 kV. Tandem mass spectrometry analysis was carried out on a Q-Exactive HFX mass spectrometer (ThermoFisher) using data-independent acquisition (DIA). DIA was performed as previously described using variable isolation widths for different m/z ranges^20^. Stepped normalized collision energy of 25 ± 10 % was used for all DIA spectral acquisitions.

Raw MS data were analyzed using quantitative DIA proteomics software DIA-NN (v. 1.8)^21^. Complete mouse proteome databased from UNIPROT was used for neural network generation, with deep spectral prediction enabled. Protease digestion was set to trypsin (fully specific) allowing for two missed cleavages and one variable modification. Oxidation of Met and acetylation of the protein N-terminus were set as variable modifications. Carbamidomethyl on Cys was set as a fixed modification. Match between runs and remove likely interferences were enabled. Neural network classifier was set to double-pass mode. Protein interferences were based on genes. Quantification strategy was set to any liquid chromatography (LC, high accuracy). Cross-run normalization was set to RT-dependent. Library profiling was set to smart profiling.

### SDS-PAGE

Protein concentrations of all fractions or insulin granule pellets from fractionation methods were analysed using a Pierce BCA Protein assay (ThermoFisher) and subjected to SDS-PAGE analysis. All primary and secondary antibodies used are listed in Table 1. Protein signals were developed and detected by chemiluminescence (Millipore) on a ChemiDoc MP Imaging System (Bio-Rad). Insulin SDS-PAGE was performed using modified SDS-PAGE protocol described previously^22^.

**Table 1.**
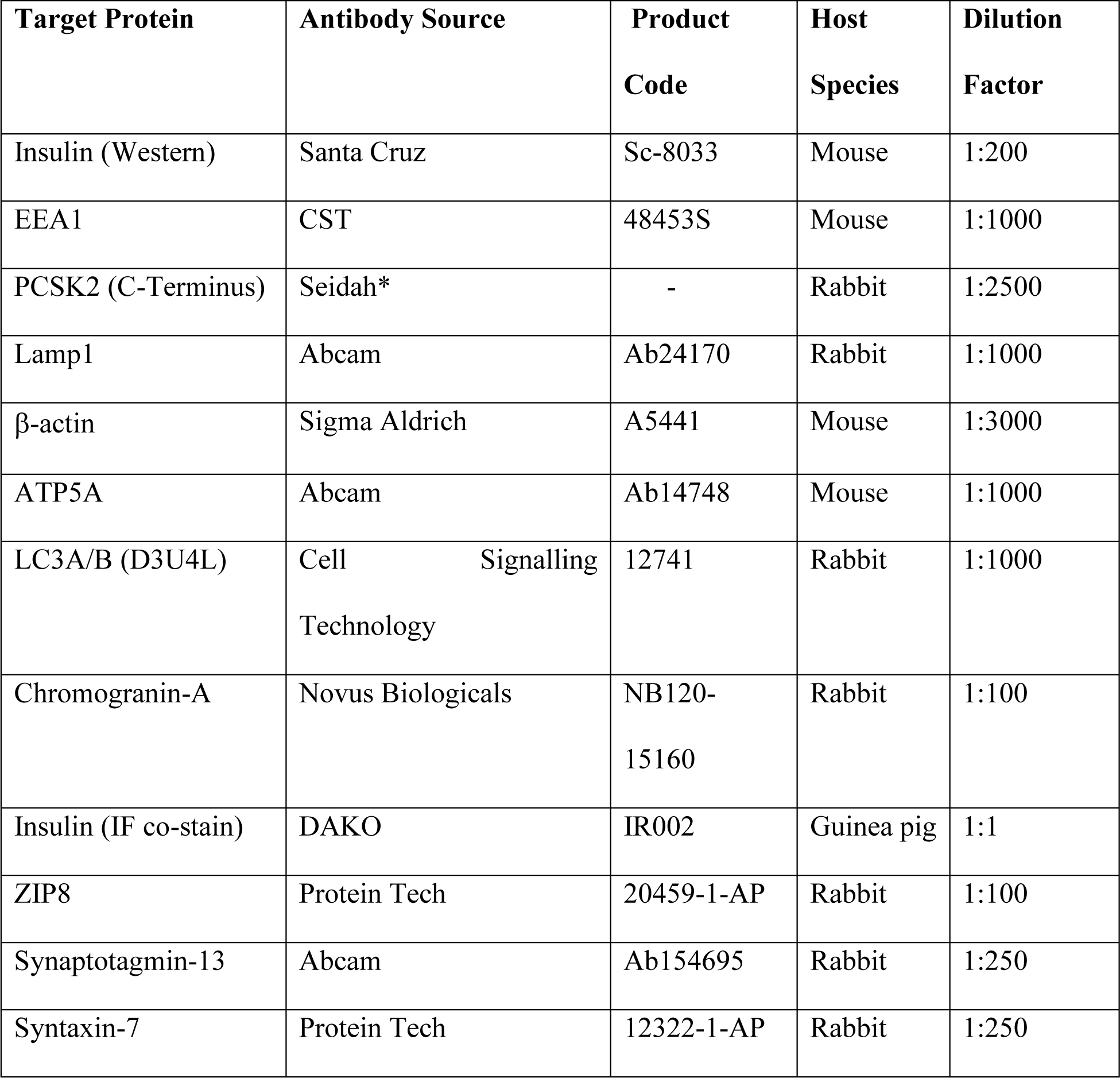
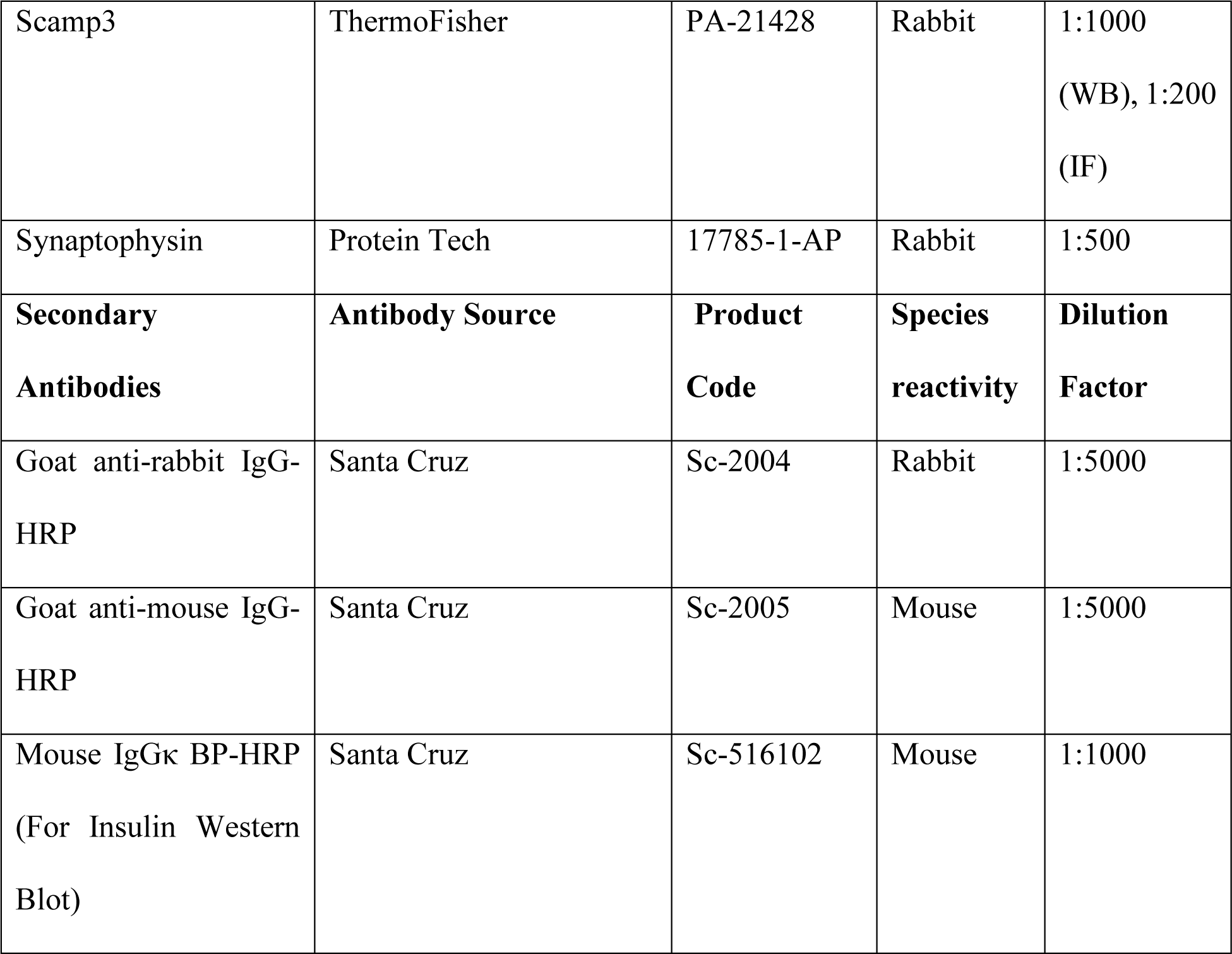
Antibodies used for Western Blot and immunofluorescent staining. *Anti-PCSK2 antibody (C-Terminus) gifted by Dr. Nabil Seidah (Clinical Research Institute of Montreal)

### Immunofluorescent Staining

Cells and human pancreata were prepared for immunofluorescent staining following standard procedure previously described^18,23^.

### Microscopy

Fixed microscope slides were imaged using the Leica LCS SP8 confocal microscope (Wetzlar, Germany). Slides were imaged using a 93X magnification glycerol lens and a white laser. Leica LAS X software was used for image collection and colocalization analysis done with Fiji ImageJ^24^.

### Insulin ELISA

Insulin concentrations were measured by insulin ELISA using commercially available Ultra-Sensitive Mouse Insulin ELISA Kit (Crystal Chem, USA) according to manufacturer’s recommendations.

### siRNA Knockdown of Scamp3

A SCAMP3 knockdown (KD) model was generated by transfecting rat insulinoma INS-1 beta-cells with siRNA mediated oligomers. INS-1 cells were cultured on glass microscope coverslips for immunofluorescent imaging or 6-well and 12-well plates for Western Blot and GSIS respectively. Cells were transfected using two commercial rat Scamp3 siRNAs (TriFECTa DsiRNA, IDs: rn.Ri.Scamp3.13.2 #613664206 (si#1), rn.Ri.Scamp3.13.3 #613664207 (si#2), and a non-targeting siRNA (TriFECTa DsiRNA, Lot #0000865210) complexed with Lipofectamine 2000 (ThermoFisher) following manufacturer protocols at a concentration of 10 nM. INS-1 cells were fixed or harvested 48 hours post transfection to validate Scamp3 KD by Western Blotting and confocal imaging. Glucose stimulated insulin secretion assays were also performed on INS-1 cells 48 hours post transfection.

### Glucose stimulated insulin secretion (GSIS)

Glucose stimulated insulin secretion in Scamp3 KD and control cells were performed as previously described^18^.

### Data analysis

All output raw MS data was analysed using Rstudio (v. 4.1.1.). Variance stabilization normalization (vsn) was performed on the data using the *vsn* package. PCP analysis was done using the functions: hclust, numClusters and cutree from Rstudio. To assign proteins to organelles and complexes within cells, we used support vector machine learning implemented with the ‘svmOptimisation’ and ‘svmClassication’ functions in the R package *pRoloc* ^25,26^. For intraclass correlation coefficients, our model fitted sample fraction IDs (Fraction 1-12) as a random effect, and normalized expression number for known ISG proteins as the outcome variable. ICC was calculated as the ratio of the between fraction variance to the total variance. To assign proteins within our samples as markers for other organelles distinct from ISGs, we used the publicly available *Mus musculus* library, available through the package *pRolocdata*^27^. GraphPad Prism (version 8.0) was used for statistical analyses. Results were considered significant at P-value < 0.05.

## RESULTS

### Optimized three-step ISG enrichment method

Our subcellular fractionation protocol uses a three-step purification of MIN6 post-nuclear cell lysates on a discontinuous gradient (8–8 - 30 %) (**Figure 1–A & B, described in detail in Materials and Methods**). Insulin concentrations were quantified in 16 Optiprep fractions, revealing two distinct peaks of insulin (**Figure 1C**) corresponding to immature ISGs (Fraction 5 – 7) and mature ISGs (Fractions 11 – 13). SDS-PAGE analyses of the fractions confirmed enrichment of mature insulin in the later fractions, and proinsulin in earlier fractions (**Figure 1D**). Furthermore, through SDS-PAGE analysis, there is a clear enrichment of mature ISGs lacking contaminants from the cytoskeleton (beta-actin) and endosomes (EEA1). Enrichment of PCSK2 (an ISG marker) was also present that matched the ISG enrichment. Fraction 13 exhibited maximal insulin enrichment but also contained contaminating mitochondria (**Figure 1D**) so Fractions 11 and 12 were chosen for subsequent purification.

**Figure 1:**
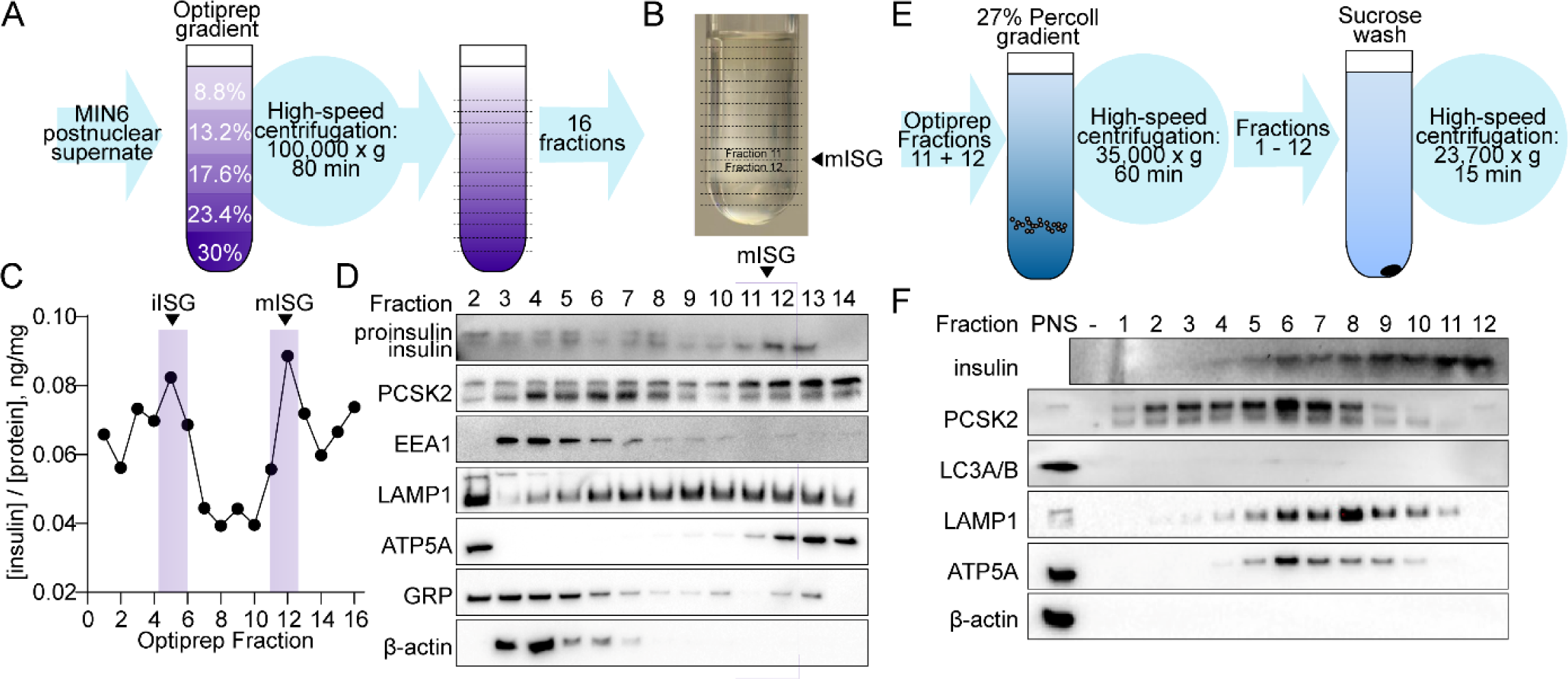
Analysis of isolation of immature and mature insulin secretory granules *from MIN6 beta-cells by Optiprep^TM^ and Percoll*. (**A**) *Optiprep workflow*: MIN6 post-nuclear supernatant is loaded atop 5 fixed concentrations of Optiprep and ultra-centrifuged for 75min at 100,000g. (**B**) Representative example of visible subcellular fractionation distribution after ultra-centrifugation. (**C**) Representative quantification of insulin enrichment from 16 fractions of Optiprep by insulin ELISA (**D**) Western blot analysis of pro- and insulin enrichment of Optiprep fractions, as well as marker proteins for subcellular components of beta-cells. (**E**) *Percoll workflow*: Fractions 11 and 12 from Optiprep gradients following insulin enrichment analysis are loaded on top of 27% Percoll and ultra-centrifuged for 60min at 35,000g. (**F**) SDS-PAGE analysis of insulin enrichment of Percoll fractionation, as well as marker proteins for subcellular components of MIN6 beta-cells.

Optiprep Fractions 11 and 12 were loaded atop a 27 % Percoll solution and ultra-centrifuged (**Figure 1E**), then washed in sucrose to enable SDS-PAGE analyses and tandem mass spectrometry. SDS-PAGE analysis using antibodies against markers of subcellular compartments for ISG (insulin, PCSK2), cytoskeleton (beta-actin), autophagosomes (LC3A/B) and mitochondria (ATP5A) (**Figure 1F**) confirmed insulin enrichment in later Percoll fractions, with minimal contaminations from other organelles removed during the first-step fractionation through Optiprep.

### Unbiased proteomics analysis of fractions from three-step purification of ISGs (PCP)

All twelve sucrose-washed fractions from each experiment underwent mass spectrometry-based proteomic analysis. Protein quantification was performed using label-free analysis in the MaxQuant package, and the protein-level data were subsequently analyzed by PCP and supervised machine learning using the R package *pRoloc* ^26,27^. Protein abundance across samples were normalized using variance stabilization normalization (VSN) using the *vsn* package^28^. Following this, the covariance of 5 known marker proteins (INS1, INS2, CPE, PCSK2, CGA) within each experiment (*n* = 7) was quantified by intraclass correlation coefficient (ICC) to evaluate experiment reliability. ICCs were calculated based on random effects using a linear mixed model implemented in the package *lme4* ^29^. Two out of seven replicate experiments exhibited low and outlying ICC values (< 0.3) and were therefore excluded from downstream analysis (**Supplementary Table 1**).

Through PCP analysis, proteins with similar abundance patterns are grouped and clustered together based on Euclidean distances are iteratively between each protein. Hierarchical clustering of all proteins (**Supplementary Table 2**) resulted in identifications of 432 significant potential candidates of ISG proteins (**Supplementary Table 3**). Of interest, a single cluster contained well-established ISG marker proteins, including Insulin (INS1 and INS2), islet amyloid polypeptide (IAPP), prohormone convertases 1 and 2 (PCSK1, PCSK22), Chromogranin A (CGA), Secretogranin-2 and −3 (SCG2, SCG3), Carboxypeptidase E (CPE), Zinc transporter 8 (SLC30A8), many proton ATP-ase subunits, syntaxins, synaptotagmins, vesicle associated membrane proteins (Vamp2, Vamp3 and Vamp7), and Rab-GTPases (Rab3a, Rab27a) (**Supplementary Table 3**), suggesting high efficiency of the three-step purification method and validity of the proteomics data. Representative protein profiles from a single replicate illustrated a high enrichment of this ISG-specific protein cluster (**Figure 2A**), further validating the purification method. Subsequent GO enrichment and STRING analysis of this ‘ISG’ cluster (**Figure 2B**) reveals a high strength index of proteins involved in insulin processing, secretory granule localization as well as granule trafficking (teal). However, some subsets of proteins within this cluster were identified as contaminants, including ribosomal subunits (yellow), lysosomal cathepsins (purple) and subunits of mitochondrial electron transport chain proteins (red). While *known* ISG proteins were identified through PCP analysis, many other proteins within this cluster may also be ISG-resident. For example, Synaptotagmin-5, and Synaptotagmin-like 4 are potential synaptotagmin isoforms present on ISGs in INS-1E cells^14^. Additionally, the identification of Synaptotagmin-13 (SYT13), Synaptophysin (SYP), zinc transporter ZIP8, and Syntaxin-7 (STX7) suggest their potential residence on ISG membranes. Other proteins of interest are protein disulfide isomerases (e.g. Pdia6, Pdia3, PDI), which have been shown to facilitate insulin processing^30^.

**Figure 2:**
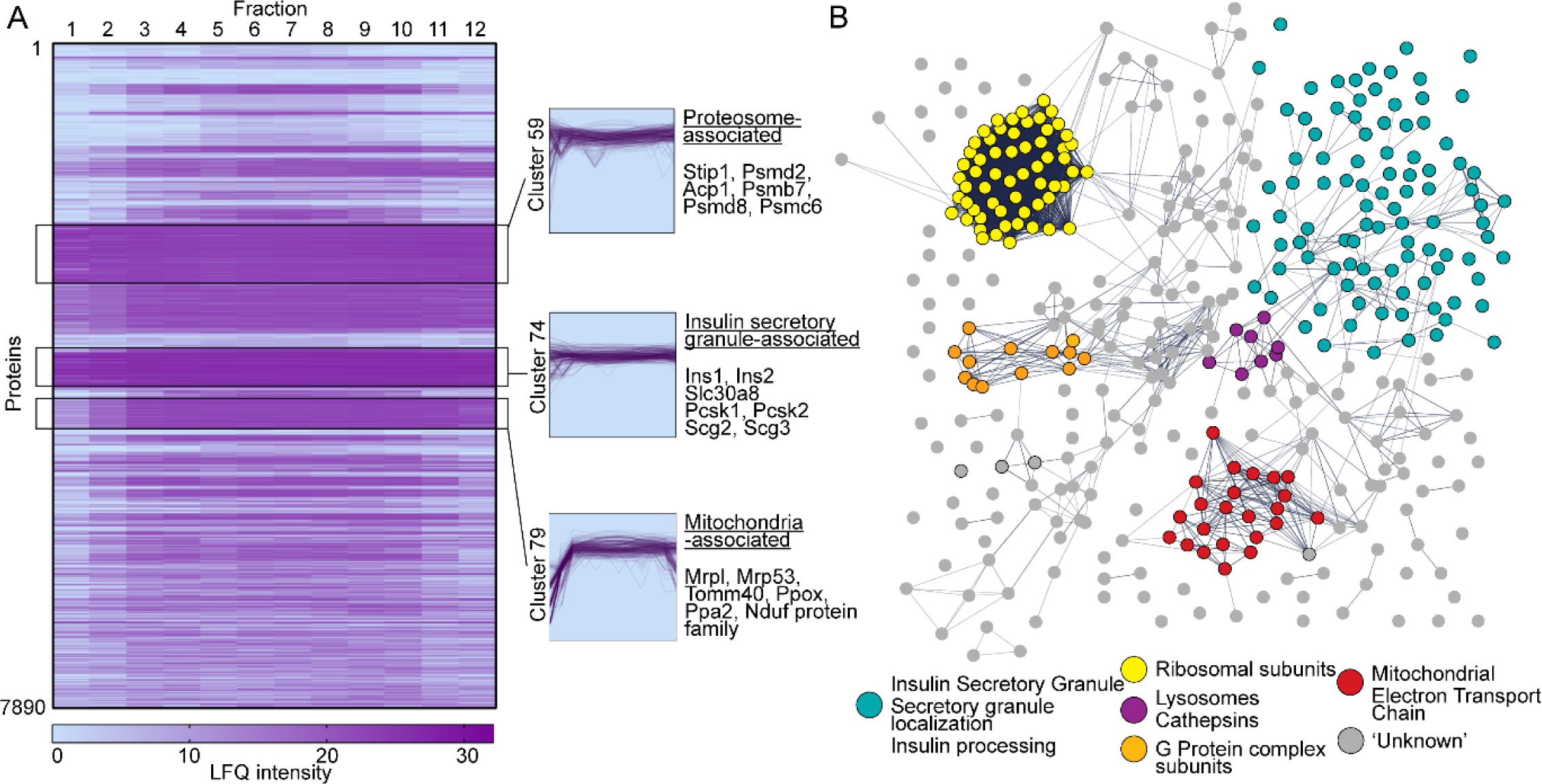
Heatmap of protein enrichment hierarchical clustering across 12 fractions *after three-step purification of mISGs*. (**A**) Representative heatmap of abundance levels of proteins across 12 fractions. All heatmaps for 5 experimental replicates (n = 5) available in **Supplementary Figure 1**. Boxes outlined indicates clusters enriched in proteasome, ISG and mitochondrial proteins. (**B**) Gene ontology enrichment analysis of Cluster 74 (ISG-associated proteins), with individual proteins coloured based on enriched biological processes.

### Targeted proteomics analysis of fractions from three-step purification of ISGs (LOPIT-DC)

PCP analyses offer comprehensive insights beyond single snapshot proteomics of ISGs, enabling unbiased matching of co-abundant proteins across subcellular fractions. While previous ISG snapshot proteomes yielded only 5 consensus proteins ^31^, our PCP analysis identified these proteins along with other well-established ISG-resident proteins: ZNT8, ProSAAS (PCSK1 precursor), IAPP, Secretogranin-2 and Secretogranin-3 ^9,32–35^. Subsequently, these 10 ISG proteins were selected as candidates for subcellular localization analyses. ‘Marker’ proteins were assigned for other non-ISG subcellular compartments of the β-cell using the *Mus musculus* library available in pRoloc. The 10 ISG proteins chosen were manually added as markers. Using a support vector machine learning (SVM; a form of supervised machine learning) within pRoloc ^26,27^, unmarked protein profiles were correlated with marker protein profiles to generate a full spatial proteome. Proteins assigned to the ISG exhibited high SVM score distributions (**Supplementary Figure 1**) across experimental replicates, indicating strong assignment accuracy. Notably, these proteins showed distinct profiles compared to proteins assigned to other subcellular localizations. With confidence, we identified a total of 211 ISG proteins across five experiment replicates (**Supplementary Table 4**). Notably, SYP, SYT13, and STX7 were among the ISG-associated proteins identified. Additionally, SCAMP3, initially missed in the PCP analysis, was found in all 5 replicates in our targeted LOPIT-DC analysis (**Supplementary Table 4**).

### Immunocytochemistry Validation of Candidate proteins

We further validated ZIP8, SYT13, STX7, SYP and SCAMP3 as ISG-resident proteins using immunocytochemistry and colocalization analyses and showed that all were present in MIN6 cells (**Figure 3A**), showing significant positive correlation with insulin in ISGs (**Figure 3B**). Chromogranin-A (CHGA), a well-established ISG cargo protein, exhibited high colocalized with insulin (**Figure 3A**), serving as a positive control. SYP was observed closer to the plasma membrane with reduced peri-nuclear distribution, while SCAMP3 localized predominately in the peri-nuclear region with sparse cytosolic distribution. ZIP8, identified solely through PCP analysis, showed lower Pearson’s correlation with insulin in MIN6 cells (**Figure 3B**), possibly explaining its absence from our machine learning approach. Furthermore, to validate the association of these proteins (excluding ZIP8) in human β-cells, we conducted immunostaining of pancreatic islets of non-diabetic human donors (**Figure 3C**). These proteins were found primarily in β-cells, demonstrating significant colocalization with insulin (**Figure 3D**), with additional staining present in other cell-types of the pancreatic islets for SYT13, SYP and SCAMP3.

**Figure 3:**
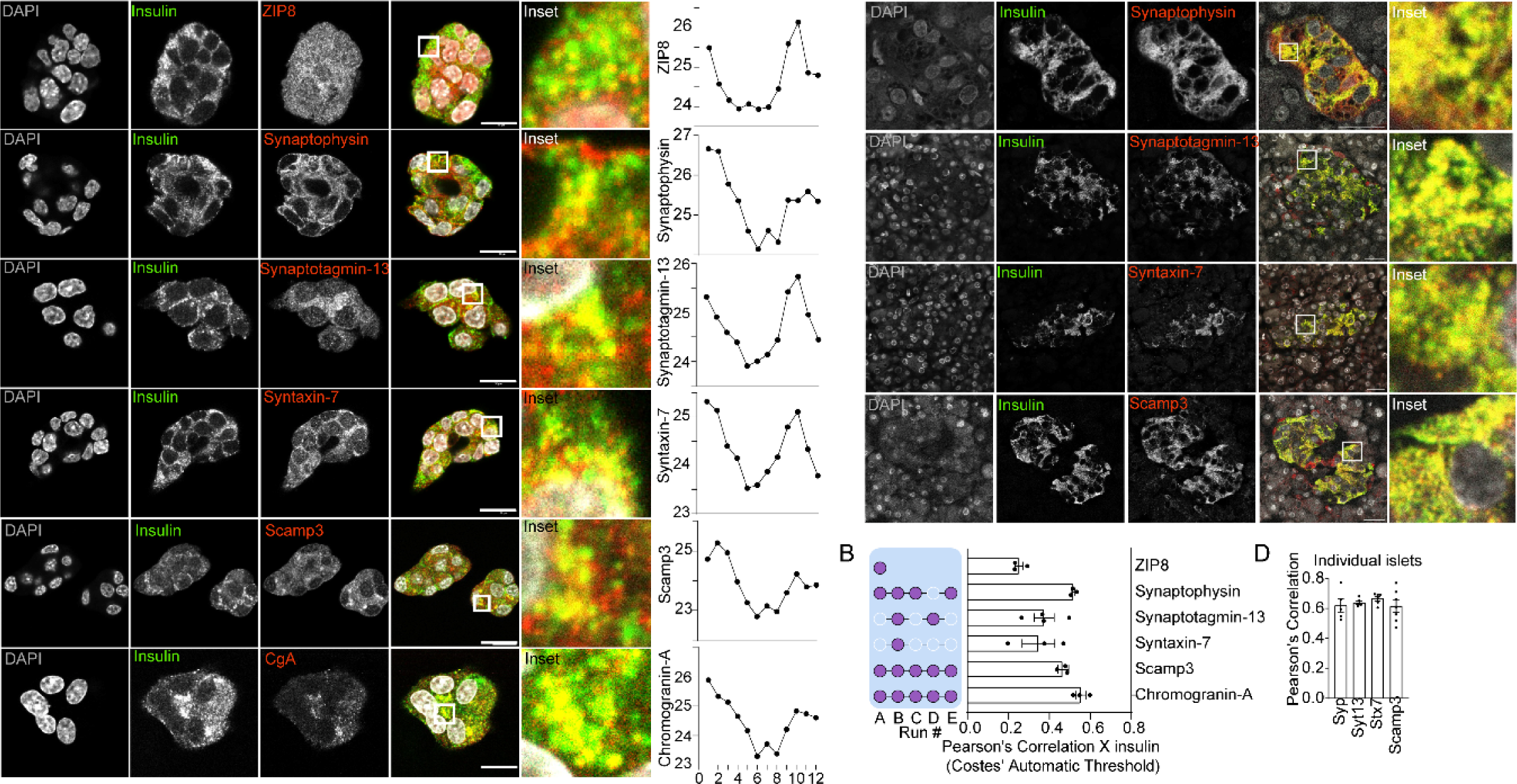
Colocalization analysis on immunofluorescent stained MIN6 cells for *candidate proteins from LC-MS/MS analysis*. (**A**) Confocal fluorescence imaging of MIN6 cells labelled with anti-insulin co-stained with anti-ZIP8 (n = 3), anti-Synaptophysin (n = 3), anti-Synaptotagmin-13 (n = 4), anti-Syntaxin-7 (n = 3), anti-Scamp3 (n = 3) and anti-Chromogranin A (n = 3). Enrichment profile of candidate proteins across 12 fractions from LC-MS/MS analysis. (**B**) Quantification of colocalization of candidate proteins with insulin using Pearson’s correlation coefficient (with Costes’ automatic thresholds). Filled dots in purple represent the number of experiments (Run #) these proteins appear within pRoloc analyses. (**C**) Confocal fluorescence imaging of human islets labelled with anti-insulin, co-stained with anti-synaptophysin (n = 1), anti-synaptotagmin-13 (n = 1), anti-syntaxin7 (n = 1) and anti-Scamp3 (n = 1). (**D**) Quantification of colocalization of candidate proteins with insulin from individual islets from a single human non-type 2 diabetic donor for each protein using Pearson’s correlation coefficient (with Costes’ automatic thresholds). All error bars represent standard error of the mean (+-S.E.M.)

### Investigation of SCAMP3 function in INS1 beta-cells

Consistent identification of SCAMP3 in all 5 experimental replicates (**Figure 3B, Supplementary Table 4**) suggested its potential role in regulating ISG-associated processes. Therefore, we chose SCAMP3 for knock-down experiments to investigate its involvement in insulin content and GSIS. Endogenous SCAMP3 in INS-1 rat insulinoma cells were depleted using commercially obtained silencer RNAs targeting two unique targeting sequences of rat SCAMP3 mRNAs (TriFECTa, IDT). At 48 hours post-transfection, INS-1 cells exhibited an ∼80% knock-down of SCAMP3 in both siRNA conditions (**Figure 4A and B**) compared to cells transfected with non-targeting control siRNA.

**Figure 4:**
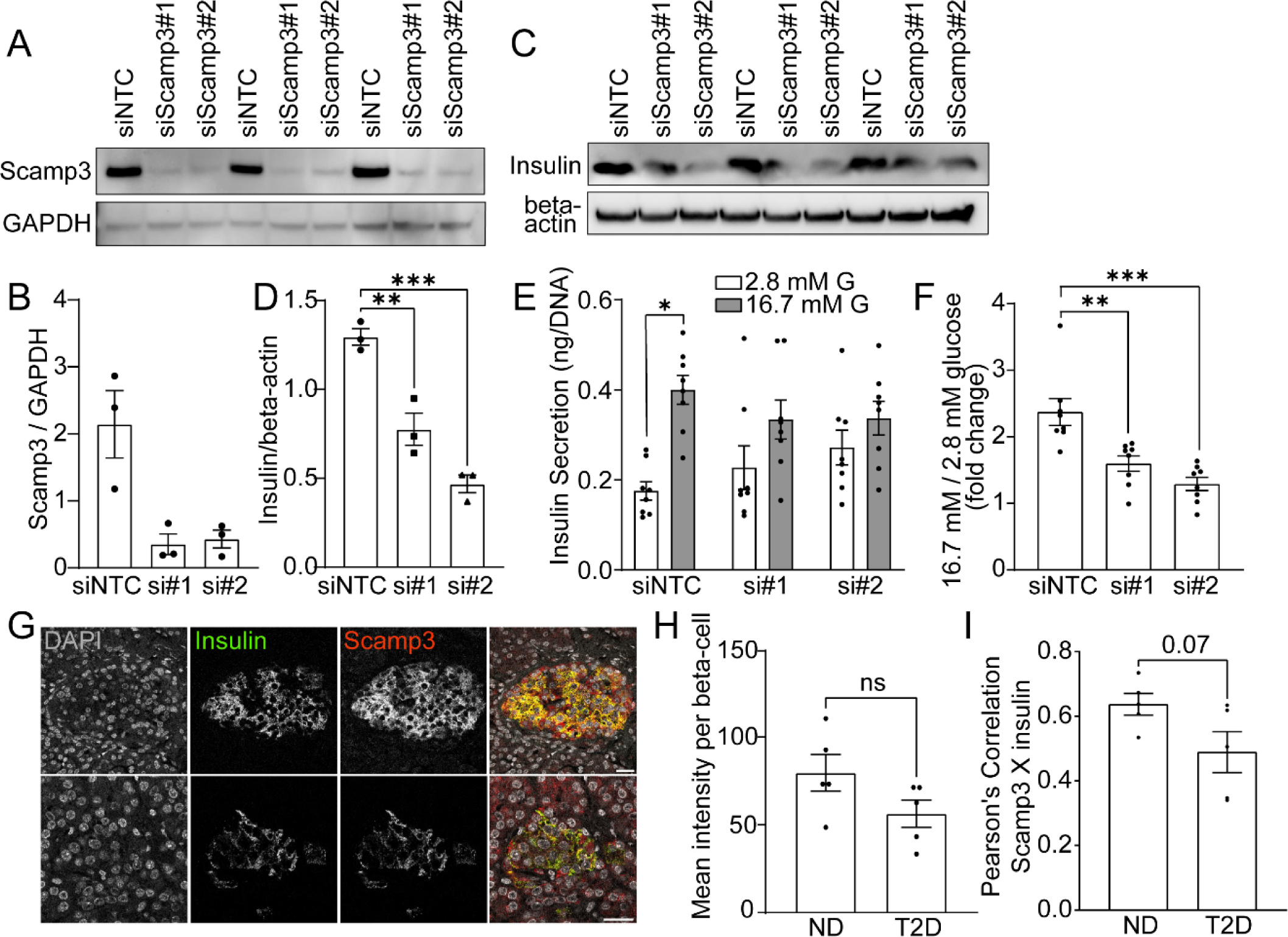
Functional investigation of Scamp3 in INS-1 β-cells and human non-type *2 diabetic and type 2 diabetic pancreatic islets*. (**A**) SDS-PAGE analysis of INS-1 cell lysates 48 hours post transfection with a control non-targeting siRNA and two unique siRNAs specific to Scamp3 (siScamp3#1 and siScamp3#2) in 3 experimental replicates together (n = 3). (**B**) Densitometry quantification of knockdown efficiency of Scamp3 48 hours post transfection (n = 3) normalized to GAPDH. (**C**) Insulin SDS-PAGE analysis of INS-1 cell lysates 48 hours post transfection of control and siRNA KD INS-1 cells. (**D**) Densitometry quantification of insulin content in control and siRNA KD INS-1 cells (n = 3) normalized to beta-actin. (**E**) Glucose-stimulated insulin secretion assay: insulin concentration after basal (2.8 mM) and stimulation (16.7 mM) glucose conditions from INS-1 control and siRNA KD cells measured by insulin HTRF (n = 8). (**F**) Fold-change of basal to stimulation glucose conditions from E (n = 8). (**G**) Representative confocal fluorescence imaging of human non-type 2 diabetic (n = 5) and type 2 diabetic (n = 5) islets labelled with anti-insulin and co-stained with Scamp3. (**H**) Mean fluorescence intensity of Scamp3 within β-cells of human non-type 2 diabetic and type 2 diabetic islets (**I**) Quantification of colocalization of Scamp3 with insulin using Pearson’s correlation coefficient (with Costes’ automatic thresholds) in human non-type 2 diabetic and type 2 diabetic islets. All error bars represent standard error of the mean (+-S.E.M.)

We found that INS1 cells depleted of SCAMP3 display reduced cellular insulin content and reduced GSIS (**Figure 4C-D and 4E**). While control cells displayed a typical insulin secretory response with a 2-fold increase in insulin secretion under the stimulated (16.7 mM) glucose condition compared to the basal (2.8 mM) glucose condition, both knock-down conditions did not exhibit significant differences between basal and stimulated conditions (**Figure 4E**). This resulted in a significant decrease in the fold changes of basal/stimulated conditions, indicating a blunted secretory response from siRNA treated cells (**Figure 4F**).

### Investigation of Scamp3 function in non-type 2 diabetic and Type 2 diabetic human pancreatic islets

To explore the role of SCAMP3 in human β-cells further, we analyzed its distribution and abundance in non-type 2 diabetic and type 2 diabetic pancreatic islets (**Figure 4G**). While a non-significant decrease in SCAMP3 fluorescence intensity was noted in type 2 diabetic islets compared to non-type 2 diabetic islets (**Figure 4H**, n=5 per group), a slight reduction in SCAMP3 and insulin localization was observed in type 2 diabetic islets (**Figure 4I**). These findings offer initial insight into potential changes in SCAMP3 expression and localization linked to type 2 diabetes.

## Discussion

Our study presents an efficient approach to isolating insulin secretory granules, successfully enriching for them using a three-step subcellular fractionation procedure. This method is now applicable to various cell lines or primary cells. Through unbiased quantitative PCP analysis, we establish the first MIN6 β-cell ISG proteome, identifying a total of 432 proteins associated with ISG. Additionally, parallel machine learning analysis enhances the proteome, refining 211 proteins assigned with high confidence to ISGs. Using these proteomes, we validate the presence of 5 ISG-resident proteins of interest in MIN6 cells, 4 of which are also validated in human donor pancreatic islets. Furthermore, through siRNA knockdown and functional analyses of SCAMP3, we validate this novel ISG-associated protein’s involvement in the glucose-regulated secretory pathway in β-cells. Expanding the use of this framework may reveal alterations in ISG morphology or proteins, enhancing our understanding of changes in granule populations in β-cell dysfunction and development of type 2 diabetes.

Previous proteomics analyses of ISGs in INS1 and INS-1E cell types have identified 50-140 candidate ISG proteins ^11–14^, with only one study using protein-correlation profiling. Here, we employ an unbiased PCP analysis to hierarchically cluster proteins with highly similar profiles, validating ISG separation techniques and identifying potential novel ISG resident proteins. Our approach identified many proteins known to be ISG localized. GO enrichment analysis of these clusters further support our findings, indicating high strength indices for insulin processing, protein localization to secretory granules, and insulin metabolic processes.

Validation is necessary for many other proteins identified through PCP analysis to investigate their potential localization to the ISG and their impacts on insulin granule processes. Specifically, from PCP alone, we chose to validate ZIP8, a metal-ion transporter. Zinc plays a crucial role in cellular signalling, and within β-cells, zinc uptake and accumulation within the ISG are critical for insulin maturation, storage of insulin and secretion ^36^. There exists a clear relationship between intracellular zinc concentrations and ISG processes. Another zinc transporter, Slc30a8 (ZnT8), is highly expressed in β-cells and localized to ISG membranes to regulate zinc concentrations within the ISG lumen ^37^. ZnT8 haploinsufficiency in MIN6 cells down-regulate mRNA levels of ZIP8 ^32^. These findings, combined with the identification of ZIP8 through PCP analysis, suggest a direct physical interaction of ZIP8 with ZnT8 on the ISG. Many other proteins from the SLC family were identified through both analyses and their validation may uncover potential novel ISG solute carriers that regulate ISG function (**Supplementary Table 3 & 4**).

Hierarchical clustering reveals the presence of many previously known ISG-associated integral membrane proteins, including members of the vSNARE family (Vamp2/3 and Vamp7), which are known contributors to insulin granule exocytosis ^38,39^. Additionally, our analysis identifies numerous Rab-GTPases including Rab3a, Rab27a, Rab8a, known small G-proteins that affect ISG movement ^40^. While Rab3a and Rab27a have established roles in ISG motility, the abundance of other Rab proteins suggests potential similar roles in ISG dynamics. Supervised machine learning analysis of the ISG proteome confirms the presence of several Rab-GTPases, including Rab8a and Rab7a.

Identified synaptic vesicle proteins such as membrane-localised synaptotagmins regulate ISGs exocytosis ^41^; Syt7 colocalizes with insulin and participates in replenishing ISGs in human islets. Syt13 in humans is abundantly expressed in the pancreas ^42^, regulates islet formation, and its knock-out affects α to β-cell ratios in islets ^43^. Syt13 is also downregulated in murine models of diabetes (*db/db*) and obesity (*ob/ob*) ^44^, demonstrating its clear involvement in β-cell development and GSIS.

Synaptotagmins also interact with syntaxin proteins (e.g. SNAP23, VAMPs) to form SNARE complexes that regulate granule fusion during insulin exocytosis in β-cells. Syntaxin-7 is identified as an ISG-associated protein and forms SNARE complexes with VAMP4, STX8 and VTI1B, facilitating proinsulin degradation within immature secretory granules ^45^. Additionally, STX7 is involved in SNARE complex formation with VAMP7 and STX8 during autophagosome formation ^46^.

Synaptophysin, typically associated with neurons and synaptic vesicle function, emerges as a novel ISG-associated protein (**Figure 3A & 3B**). While its roles in β-cells, possibly involving GABA secretion via synaptic-like microvesicles is debated ^47^, we have observed its upregulation in murine islets under conditions of high-fat diet and in mouse models of diabetes ^44^. Our findings show the precise role of synaptophysin in β-cells warrants further investigation.

### Identification of novel ISG protein Scamp3

One of the most intriguing ISG-associated proteins we uncovered is SCAMP3. While SCAMP protein family is widely expressed and found on post-Golgi vesicle membranes ^48^, SCAMP3’s role in β-cells remained elusive. However, other SCAMP proteins have been implicated in GSIS. Our study reveals clear association of SCAMP3 within ISGs in rat, mouse, and human β-cells, and knocking down SCAMP3 significantly reduces GSIS (**Figure 4F**) and insulin content (**Figure 4D**). A single transcriptome-wide association study has also shown the presence of splice sites in the SCAMP3 gene that associates with type 2 diabetes susceptibility in human pancreatic islets ^49^. In addition, previous proteomics studies have shown that mouse pancreatic islets exposed to high glucose result in a downregulation of SCAMP3 compared to islets exposed to low glucose conditions ^50^. In summary, our data provide the first evidence of a role of SCAMP3 in the ISG biogenesis, trafficking, or secretory pathway. Further studies should aim to further understand the precise mechanisms by which SCAMP3 regulates insulin secretion and insulin stores in the β-cell.

## Acknowledgments

We thank the core facilities at the Charles Perkins Centre at the University of Sydney. This study was made possible with access to the Sydney Mass Spectrometry and Sydney Microscopy & Microanalysis facilities. We thank the organ donors and their families for their generosity. We also thank the members of St Vincent’s Institute in the islet isolation program and Donatelife for providing research consent and provision of human pancreata. St Vincent’s Institute receives support from the Operational Infrastructure Support Scheme of the Government of Victoria. This work was supported by National Health and Medical Research Council (NHMRC) project grant GNT1139828 (to M.A.K.). N.N is supported by the University of Sydney Postgraduate Award (UPA).

## Resource Availability

### Corresponding author

Further information requests for reagents and resources used should be directed to lead contact, Melkam Kebede.

### Materials availability

This study did not generate any new unique reagents.

### Data code availability

Any additional information required to re-analyze the data reported in this paper is available upon request from the lead contact. This paper does not produce original code. The mass spectrometry proteomics data have been deposited to the ProteomeXchange Consortium via the PRIDE partner repository with the dataset identifier PXD051136, Username, Password: GnZTdMqQ”.

### Author Contributions

Conceptualization, M.A.K. and B.Y.; Methodology, N.N. and B.Y.; Validation, M.A.K. and B.Y.; Formal Analysis, N.N., A.M.S. and M.L.; Investigation, N.N., B.Y., C.F. and M.L.; Resources, M.A.K., Data Curation, N.N., B.Y., M.L and A.M.S; Writing – Original Draft, N.N.; Writing – Review & Editing, M.A.K., B.Y., M.A.S. and M.L.; Visualization, B.Y.; Supervision, M.A.K.; Project Administration, M.A.K. and B.Y.; Funding Acquisition, M.A.K.

